# SynLight: a dicistronic strategy for simultaneous active zone and cell labeling in the *Drosophila* nervous system

**DOI:** 10.1101/2023.07.17.549367

**Authors:** Michael A. Aimino, Jesse Humenik, Michael J. Parisi, Juan Carlos Duhart, Timothy J. Mosca

## Abstract

At synapses, chemical neurotransmission mediates the exchange of information between neurons, leading to complex movement behaviors and stimulus processing. The immense number and variety of neurons within the nervous system makes discerning individual neuron populations difficult, necessitating the development of advanced neuronal labeling techniques. In *Drosophila*, Bruchpilot-Short and mCD8-GFP, which label presynaptic active zones and neuronal membranes, respectively, have been widely used to study synapse development and organization. This labeling is often achieved via expression of two independent constructs by a single binary expression system, but expression can weaken when multiple transgenes are expressed by a single driver. Ensuring adequate expression of each transgene is essential to enable more complex experiments; as such, work has sought to circumvent these drawbacks by developing methods that encode multiple proteins from a single transcript. Self-cleaving peptides, specifically 2A peptides, have emerged as effective sequences for accomplishing this task. We leveraged 2A ribosomal skipping peptides to engineer a construct that produces both Bruchpilot-Short and mCD8-GFP from the same mRNA, which we named SynLight. Using SynLight, we visualized the putative synaptic active zones and membranes of multiple classes of olfactory, visual, and motor neurons and observed correct separation of signal, confirming that both proteins are being generated separately. Furthermore, we demonstrate proof-of-principle by quantifying synaptic puncta number and neurite volume in olfactory neurons and finding no difference between the synapse densities of neurons expressing SynLight or neurons expressing both transgenes separately. At the neuromuscular junction, we determined that synaptic puncta number labeled by SynLight was comparable to endogenous puncta labeled by antibody staining. Overall, SynLight is a versatile tool for examining synapse density in any nervous system region of interest and allows new questions to be answered about synaptic development and organization.

## INTRODUCTION

Synapses in the brain facilitate the exchange of information from one neuron to another, culminating in integrated signals that inform the complex computations underlying movement and stimulus sensation. The presynaptic side of the synapse is defined by the active zone, a structural site comprised of quantal release machinery that anchors calcium channels near synaptic vesicles containing neurotransmitter (Ehmann et al., 2018; Südhof, 2012; Wagh et al., 2006). After an influx of calcium through active zone-associated channels following an action potential, neurotransmitter is released into the synaptic cleft where it binds to cognate neurotransmitter receptors on the postsynaptic membrane (de Ramon Francàs et al., 2017; Lin & Goodman, 1994; Mosca & Luo, 2014; Siddiqui & Craig, 2011; Wilson, 2013). The activation of postsynaptic receptors propagates the signal from the presynaptic neuron to the postsynaptic cell. In the absence of active zones, synaptic communication is drastically impaired or blocked, and function is abrogated. Therefore, the specialized sites of communication between neurons must develop correctly over a specific timeframe and maintain their precise organization throughout an organism’s life to ensure that synaptic function continues unabated and retains aspects of synaptic plasticity necessary for appropriate behavioral coordination (Aimino et al., 2023; Farhy-Tselnicker & Allen, 2018; Silbereis et al., 2016; Waites et al., 2005). Defects in the development of active zones (both in number and organization) have been found to underlie neurodevelopmental, neuropsychiatric, and even neurodegenerative diseases including autism and schizophrenia, demonstrating the necessity for understanding how synapses develop and organize at the circuit level (Bennett, 2011; Bonansco & Fuenzalida, 2016; Grant, 2012; Mullins et al., 2016).

In the central nervous system especially, the high density of synaptic connections and concomitant difficulties of discerning the specific cell or cell-type to which an active zone localizes makes the study of synapses with cell-type specific resolution a challenge. As such, studies using antibodies to active zone machinery are limited in their utility for asking cell-type specific questions as they recognize synapses in all cells that express active zone proteins in vivo. To better discern how synapses develop and change over time with cell-type specificity, genetic strategies using binary expression systems and transgenic active zone labels have become powerful for studying synaptic organization in identified cell types with high resolution. Work in *Drosophila* especially has contributed greatly to the study of synaptic organization due to the wide variety of genetic tools available for experimental use in synaptic and neuronal labeling (Duhart & Mosca, 2022). In recent years, our understanding of synaptic biology has markedly advanced through the study of the active zone protein Bruchpilot, the orthologue of vertebrate ELKS/CAST (Ohtsuka et al., 2002), which is an essential presynaptic component at both peripheral and central synapses in the fly (Fouquet et al., 2009; Ohtsuka et al., 2002; Südhof, 2012; Wagh et al., 2006; Wang et al., 2002). In particular, the transgenic construct, Bruchpilot-Short (Fouquet et al., 2009), a truncated version of Bruchpilot that co-localizes with endogenous full-length Bruchpilot, is frequently used to study synaptic organization in specific cell-types via binary expression systems like UAS/GAL4 (Aimino et al., 2023; Berger-Müller et al., 2013; Coates et al., 2017, 2020; Duhart & Mosca, 2022; Kremer et al., 2010; Mosca et al., 2017; Mosca & Luo, 2014; Sugie et al., 2015). When Bruchpilot-Short is conjugated to a fluorescent tag and visualized using confocal microscopy, the construct appears as puncta that can be quantified, acting as a proxy measurement for the number of synapses within a specific brain region (Mosca & Luo, 2014). Use of Bruchpilot-Short has led to novel discoveries about the development and organization of synapses (Aimino et al., 2023; Berger-Müller et al., 2013; Christiansen et al., 2011; Coates et al., 2017, 2020; Fouquet et al., 2009; Kremer et al., 2010; Mosca et al., 2017; Mosca & Luo, 2014; Parisi et al., 2023), demonstrating its effectiveness as a reagent. Bruchpilot-Short is frequently used in conjunction with other synaptic and cellular transgenic constructs to label other synaptic components, including synaptic vesicles (Certel, McCabe, et al., 2022; Certel, Ruchti, et al., 2022; Estes et al., 2000; Williams et al., 2019; Zhang et al., 2002), other active zone components (Fouquet et al., 2009; Fulterer et al., 2018; Liu et al., 2011; Mosca et al., 2017; Özel et al., 2019), general cellular architecture (Lee & Luo, 1999; Potter et al., 2010; Venken, Simpson, et al., 2011), and post-synaptic sites (Andlauer et al., 2014; Fendl et al., 2020; Kremer et al., 2010; Leiss, Groh, et al., 2009; Leiss, Koper, et al., 2009; Mosca et al., 2017; Mosca & Luo, 2014; Nicolaï et al., 2010; Parisi et al., 2023; Sánchez-Soriano et al., 2005). One such construct, mCD8-GFP (T. Lee et al., 1999), is a membrane-bound GFP tag that, when expressed in any cell-type, labels the membranes of that cell, revealing its architecture. When expressed in a particular population of neurons under the control of a binary expression system, mCD8-GFP serves as a general neurite label for both dendrites and axons. Just as Bruchpilot-Short puncta can be quantified as a proxy for the number of synaptic contacts (Aimino et al., 2023; Berger-Müller et al., 2013; Christiansen et al., 2011; Coates et al., 2017, 2020; Fouquet et al., 2009; Kremer et al., 2010; Mosca et al., 2017; Mosca & Luo, 2014; Parisi et al., 2023), membrane markers like mCD8-GFP can be quantified to determine the total volume of neurite membrane in a defined population of neurons (Aimino et al., 2023; Mosca et al., 2017; Mosca & Luo, 2014). Together with Bruchpilot-Short, quantification of puncta number and neurite volume can be expressed as synaptic density within that neuronal population (Aimino et al., 2023; Mosca et al., 2017; Mosca & Luo, 2014). Thus, employing constructs like Burchpilot-Short and mCD8-GFP together makes it possible to ask vital questions about the formation and organization of synapses within a defined population of neurons, thus overcoming previous challenges brought about by the density of synaptic populations in vivo in the central nervous system.

To quantify metrics like synaptic density, it is necessary to express multiple effector or reporter constructs in vivo in tandem as synaptic density is the measure of the number of synaptic puncta within a given volume of neuronal membrane. While increasing the number of simultaneously expressed transgenes enables more complex experimentation, it also carries drawbacks. In cultured cells, multiple plasmids must be co-transfected, potentially resulting in cells that do not incorporate every plasmid that is transfected (González et al., 2011). Similarly, driving expression of multiple genes with a binary expression system like GAL4/UAS can lead to a dilution of transgene expression, causing effectors and reporters to not function optimally or sufficiently (Brand & Perrimon, 1993). While it is possible to recombine some transgenes onto the same chromosome, this is often time-consuming and challenging, depending on the chromosomal position of each transgene. Further, though it may enable the introduction of additional transgenes and reduce the difficulty of the genetic crosses needed to create the experimental animal, recombination does not reduce the genetic load of the system. Additionally, viral technologies including AAV-based vectors are often limited by the amount of genetic material that can be introduced inside a single viral particle (Grieger & Samulski, 2005; Z. Wu et al., 2010). As a result, strategies are needed to reduce the size of genetic material introduced but also to increase the likelihood of introducing all of the desired transgenic components with a reduced risk of dilution and failed or reduced expression. One approach to mitigating such drawbacks is to express a single open reading frame that can encode multiple gene products. Initially, internal ribosome entry site (IRES) sequences were used to promote the internal initiation of translation of two separate proteins (Douin et al., 2004; Martínez-Salas, 1999). However, these sequences were limited in their effectiveness as the peptide following the IRES sequence would often have decreased expression compared to the preceding peptide (Kaufman et al., 1991; Ye et al., 1997). More recent work employs 2A viral peptides, which are highly efficient ribosomal skipping peptides (Daniels et al., 2014; Diao & White, 2012; Kim et al., 2011). When incorporated into a particular mRNA, the 2A peptide sequence serves as a skipping site, allowing the ribosome to separate from the mRNA, completing translation of the first sequence, and then re-enter the mRNA at the beginning of the second sequence, starting translation anew and producing a second product. As a result, placing the sequence of a 2A peptide between two complete sequences enables one continuous mRNA to code for two or more polypeptides of interest from a single promoter (Daniels et al., 2014; Kim et al., 2011; Tang et al., 2009). This can decrease the genetic load on a particular system, as now only one transgene ensures the expression of multiple products.

In *Drosophila*, there is a multitude of binary expression system drivers (GAL4, lexA, QF) available, enabling tissue-specific expression in nearly any desired nervous system region and cell type. However, there is considerable variability in the expression strength of binary expression system driver lines, such that not all driver lines can enable transgene expression equivalently. Some cell-type specific GAL4 lines are weakly expressing, making it difficult to express multiple UAS transgenes simultaneously for concurrent labeling of multiple targets. To overcome this expression issue and enable expression of multiple neural markers via a single transgenic construct, we used a vector that contains the 2A peptide from the porcine teshchovirus-1 (P2A) coding sequence (Daniels et al., 2014) to create a single transgenic construct that encodes both mCD8-GFP and Bruchpilot-Short-mStraw from the same coding sequence. This new construct makes it possible to express both neuronal labels at the same time while simultaneously reducing the genetic load on the system. Therefore, even weakly expressing GAL4s would be able to drive expression of both proteins with high signal fidelity. As the independent constructs fluorescently label synapses as well as the neuronal membrane for visualization at the level of light microscopy, we have named the tool SynLight, a portmanteau of synapse and light. We designed versions of SynLight that can be driven via multiple binary expression systems, including the GAL4/UAS and QF/QUAS systems, making it possible to ask new questions about the development and organization of synapses throughout the nervous system with less concern over transgenic dilution or failed labeling. Here, we validate SynLight expression in multiple regions of the adult and larval nervous systems in *Drosophila*, including the olfactory, visual, and neuromuscular systems using both the GAL4/UAS and QF/QUAS systems. We additionally demonstrate that SynLight expression does not affect normal neuronal morphology nor active zone puncta number as measurements from SynLight expression are quantitatively indistinguishable from measurements via independent Bruchpilot-Short-mStraw or mCD8-GFP transgene expression as well as endogenous Bruchpilot antibody staining. Thus, SynLight labels presynapses and neurite membranes, facilitating their visualization with high resolution and permitting more complex experimental design with reliable quantitative measurements. When expressed in a cell-type specific manner using existing GAL4 or QF promoter lines, SynLight has a wide applicability to a variety of *Drosophila* nervous system regions and is a versatile tool for studying synaptic development and organization.

## METHODS AND MATERIALS

### Fly stocks and care

All control lines and genetic fly stocks were maintained on cornmeal::dextrose medium (Archon Scientific, Durham, NC) at 21°C while crosses were raised on similar medium at 25°C (unless noted in the text) in incubators (Darwin Chambers, St. Louis, MO) at 60% relative humidity with a 12/12 light/dark cycle.

Transgenes were maintained over balancers with fluorescent markers and visible phenotypic traits to allow for selection of adults and larvae of the desired genotype. To drive expression in specific classes of CNS neurons, we used the following GAL4 or QF expression lines: Or47b-GAL4 (Vosshall et al., 2000), Or67d-GAL4 (Stockinger et al., 2005), Or67d-QF (Liang et al., 2013), Mz19-GAL4 (Jefferis et al., 2003), NP3056-GAL4 (Y. H. Chou et al., 2010), DIP-γ-GAL4 (Carrillo et al., 2015), and 27B03-GAL4 (Jenett et al., 2012). Expression at the NMJ was achieved via elavC155-GAL4 (Lin & Goodman, 1994) and n-syb-QF (Riabinina et al., 2015). The following UAS transgenes were used as synaptic labels or to express molecular constructs for genetic perturbation experiments: UAS-Brp-Short-mStraw (Fouquet et al., 2009; Mosca & Luo, 2014), UAS-mCD8-GFP (T. Lee & Luo, 1999), UAS-SynLight (UAS-mCD8-GFP-P2A-Brp-Short-mStraw, this study).

### Cloning of SynLight Plasmid and Transgenic Lines

Using restriction enzyme cloning, we first inserted the mCD8-GFP sequence (from pC-attB-bursα-mCD8-GFP-T2A-GAL4; a gift from Benjamin White) into a plasmid containing pC5-P2A-KAN (Daniels et al., 2014). We used BamHI and Stul as cut sites to put this sequence upstream of the P2A peptide sequence. We subsequently inserted the Bruchpilot-Short-mStrawberry sequence (pENTR-Brp-Short-mStraw; Mosca & Luo, 2014), using Nhel and AvrII as cut sites to put the sequence downstream of the P2A peptide. This strategy kept all sequences in frame. mCD8-GFP-P2A-Bruchpilot-Short-mStrawberry was then migrated from the shuttle vector into a plasmid containing a pUAS-C5-attB sequence using restriction enzyme cloning with Fsel and AscI as the cut sites and ligated together to produce the final plasmid. A similar approach was used to engineer the QUAS version of the plasmid (pQUAST-Brp-Short-mStraw-attB; Mosca & Luo, 2014). The final plasmid was sequence-verified (GeneWiz, South Plainfield NJ) and the final construct sequence is available upon request. A Qiagen Maxi Prep kit (Qiagen, cat. no. 12163) was used to isolate donor plasmid DNA for creation of transgenic fly lines. Transgenic flies (UAS and QUAS lines) were generated (BestGene, Chino Hills, CA) with the construct integrated into the attP2 docking site (Groth et al., 2004) on the 3rd chromosome. Subsequent transgenic flies (UAS line) were also generated (BestGene, Chino Hills, CA) with the construct integrated into the VK00037 docking site (Venken et al., 2006) on the 2nd chromosome.

### Immunocytochemistry

Adult flies were cleared from vials one day before collection and on the following day, newly eclosed adults were chosen based on genotype using identifiable balancers and phenotypic markers. Flies were then aged ten days before dissection and immunostaining. Brains were fixed in 4% paraformaldehyde for 20 minutes before being washed in phosphate buffer (1x PB) with 0.3% Triton (PBT). Brains were then blocked for an hour in PBT containing 5% normal goat serum (NGS) before being incubated in primary antibodies diluted in PBT with 5% NGS for two days at 4°C. Following staining, primary antibodies were discarded and the brains washed 3 × 20’ with PBT and incubated in secondary antibodies diluted in PBT with 5% NGS for an additional two days at 4°C. The secondary antibodies were then discarded, the brains washed 3 × 20’ in PBT, and then incubated overnight in SlowFade™ (ThermoFisher Scientific, Waltham, MA) gold antifade mounting media and allowed to sink. Brains were then mounted in SlowFade mounting media using a bridge-mount method with No. 1 cover glass shards and stored at 20°C before being imaged (J. S. Wu & Luo, 2006).

Larvae were processed for immunocytochemistry as previously described (Mosca & Schwarz, 2010; Restrepo et al., 2022). Larvae were grown in population cages on grape plates with yeast paste until they reached wandering third instar stage. Larval fillet dissections were done in Ca^2+^-free modified *Drosophila* saline and then fillets were fixed in 4% paraformaldehyde in phosphate buffered saline (PBS) for 20 minutes. Samples were then washed with phosphate buffered saline with 0.3% Triton (PBST). The fillets were blocked with PBST containing 5% NGS for 1 hour at room temperature and then incubated in primary antibodies diluted in PBST with 5% NGS overnight at 4°C. The following day, primary antibodies were discarded and then fillets were washed with PBST before being placed in secondary antibodies diluted in PBST with 5% NGS for 2 hours at room temperature. Fillets were mounted in Vectashield (Vector Laboratories) and stored at 4°C before being imaged.

The following primary antibodies were used: mouse anti-Nc82 (DSHB, cat. no. mAbnc-82, 1:250; Laissue et al., 1999), rabbit anti-DsRed (TaKaRa Bio, cat. no. 632496, 1:250; Mosca & Luo, 2014), chicken anti-GFP (Aves, cat. no. GFP-1020, 1:1000; Mosca & Luo, 2014), rat anti-N-Cadherin (DSHB, cat. no. mAbDNEX-8, 1:40; Hummel & Zipursky, 2004), and Alexa647-conjugated goat anti-HRP (Jackson ImmunoResearch, cat. no. 123-605-021, 1:100; Jan & Jan, 1982). Alexa488-(Jackson ImmunoResearch, West Grove, PA), Alexa568-(ThermoFisher Scientific, Waltham, MA), and Alexa647-conjugated (Jackson ImmunoResearch, West Grove, PA) secondary antibodies were used at 1:250 while FITC-conjugated (Jackson ImmunoResearch, West Grove, PA) secondary antibodies were used at 1:200. In some cases, non-specific background recognized by the dsRed antibodies (in the form of large red spots appear around the antennal lobes and outside of the tissue observed). These are part of the background, are not caused by any of the transgenic constructs used (Mosca & Luo, 2014) and did not influence any quantification or scoring methods (see below).

### Imaging and Analysis

All images of adult brains were obtained using a Zeiss LSM880 Laser Scanning Confocal Microscope (Carl Zeiss, Oberlochen, Germany) using a 20X 0.8 NA Plan-Apochromat lens, 40X 1.4 NA Plan-Apochromat lens, or a 63X 1.4 NA Plan-Apochromat f/ELYRA lens at an optical zoom of 3x. Images of third instar larval NMJs were obtained using the same confocal microscope using a 40X 1.4 NA Plan-Apochromat lens or a 63X 1.4 NA Plan-Apochromat f/ ELYRA lens. Images were centered on the glomerulus or NMJs of interest and the z-boundaries were set based on the appearance of the synaptic labels, Brp-Short-mStraw or mCD8-GFP. Images were analyzed three dimensionally using the Imaris Software 9.7.1 (Oxford Instruments, Abingdon, UK) on a custom-built image processing computer (Digital Storm, Fremont, CA) following previously established methods (Aimino et al., 2023). For both adult brains and larval NMJs, Brp-Short and endogenous Brp puncta were quantified using the “Spots” function with a spot size of 0.6 μm. Neurite volume was quantified using the “Surfaces” function with a local contrast of 3 μm and smoothing of 0.2 μm for Or47b ORNs. The resultant masks were then visually inspected to ensure their conformation to immunostaining.

### Quantitative Measurement and Statistical Analyses

All data was analyzed using Prism 8 (GraphPad Software, Inc., La Jolla, CA). This software was also used to generate graphical representations of data. Unpaired Student’s t-tests were used to determine significance between two groups while paired Student’s t-tests were used to determine significance between puncta number for individual NMJs. A p-value of 0.05 was set as the threshold for significance in all studies. For each figure, informative genotypes have been presented along with controls appropriate for each genotype.

## RESULTS

### SynLight is designed to label active zones and neurite membranes in the same cell population

Established approaches in *Drosophila* to examine synapse density in particular classes of neurons typically involve expressing two constructs: 1) Bruchpilot-Short-mStrawberry (Brp-Short-mStraw) to label active zones made by the neurons and 2) mCD8-GFP (or an equivalent) to label the processes of neurons (Aimino et al., 2023; Berger-Müller et al., 2013; Christiansen et al., 2011; Coates et al., 2017, 2020; Fouquet et al., 2009; Kremer et al., 2010; Mosca et al., 2017; Mosca & Luo, 2014; Parisi et al., 2023; Sugie et al., 2015). Previous work established that 2A peptide approaches work efficiently in *Drosophila* for encoding multiple proteins from a single ORF (Daniels et al., 2014; Diao & White, 2012). Therefore, we designed and built a single construct containing Bruchpilot-Short-mStrawberry and mCD8-GFP separated by a 2A peptide and named it SynLight. SynLight takes advantage of P2A, a viral 2A peptide from porcine teschovirus-1 (Daniels et al., 2014; Kim et al., 2011). The P2A protein is derived from a ribosomal skipping protein that allows multiple separate proteins to be made from a single mRNA (Kim et al., 2011; Tang et al., 2009). Using a vector that contains the P2A coding sequence, multiple cloning sites, and restriction sites (Daniels et al., 2014; Le et al., 2007), we engineered a single transgene that produces multiple proteins from a single coding sequence via restriction cloning (Figure 1A). The resultant vector contained mCD8-GFP inserted into the first restriction site and Brp-Short-mStraw inserted into the second site with the two separated by P2A, producing mCD8-GFP-P2A-Brp-Short-mStraw (Figure 1B). From the single mRNA produced by the transgene following activation by a promoter driver line, two separate proteins would be produced, Brp-Short-mStraw and mCD8-GFP (Figure 1C). We engineered both UAS and QUAS versions of SynLight and established transgenic lines on the 3rd chromosome in the attP2 site (Groth et al., 2004) so that the construct could be used with multiple binary expression systems. We subsequently established a transgenic line on the 2nd chromosome in the VK00037 site (Venken et al., 2006). When SynLight was expressed in olfactory neurons of the adult brain using NP3056-GAL4 (Figure 1D-D”; Y. H. Chou et al., 2010) after immunohistochemical staining for Brp-Short-mStraw and mCD8-GFP as established previously (Aimino et al., 2023; Mosca et al., 2017; Mosca & Luo, 2014), we observed clear, subcellularly distinct signal for both Brp-Short and mCD8-GFP. We obtained similar findings when SynLight was expressed in the ventral nerve cord of third instar larvae using the pan-neuronal QF driver n-syb-QF and visualized using the native fluorescence of both labels (Figure 1E-E”; Riabinina et al., 2015). For both the central and peripheral nervous systems, these data demonstrated that there was separation between the two products and indicated that ribosomal skipping occurred successfully during translation, resulting in the synthesis of Brp-Short and mCD8-GFP separately (insufficient separation would manifest as precise overlap between the mCD8-GFP and Brp-Short-mStraw channels). With the successful establishment of transgenic UAS-and QUAS-SynLight lines, we further sought to validate the construct as a synaptic labeling and quantification method.

**Figure 1.**
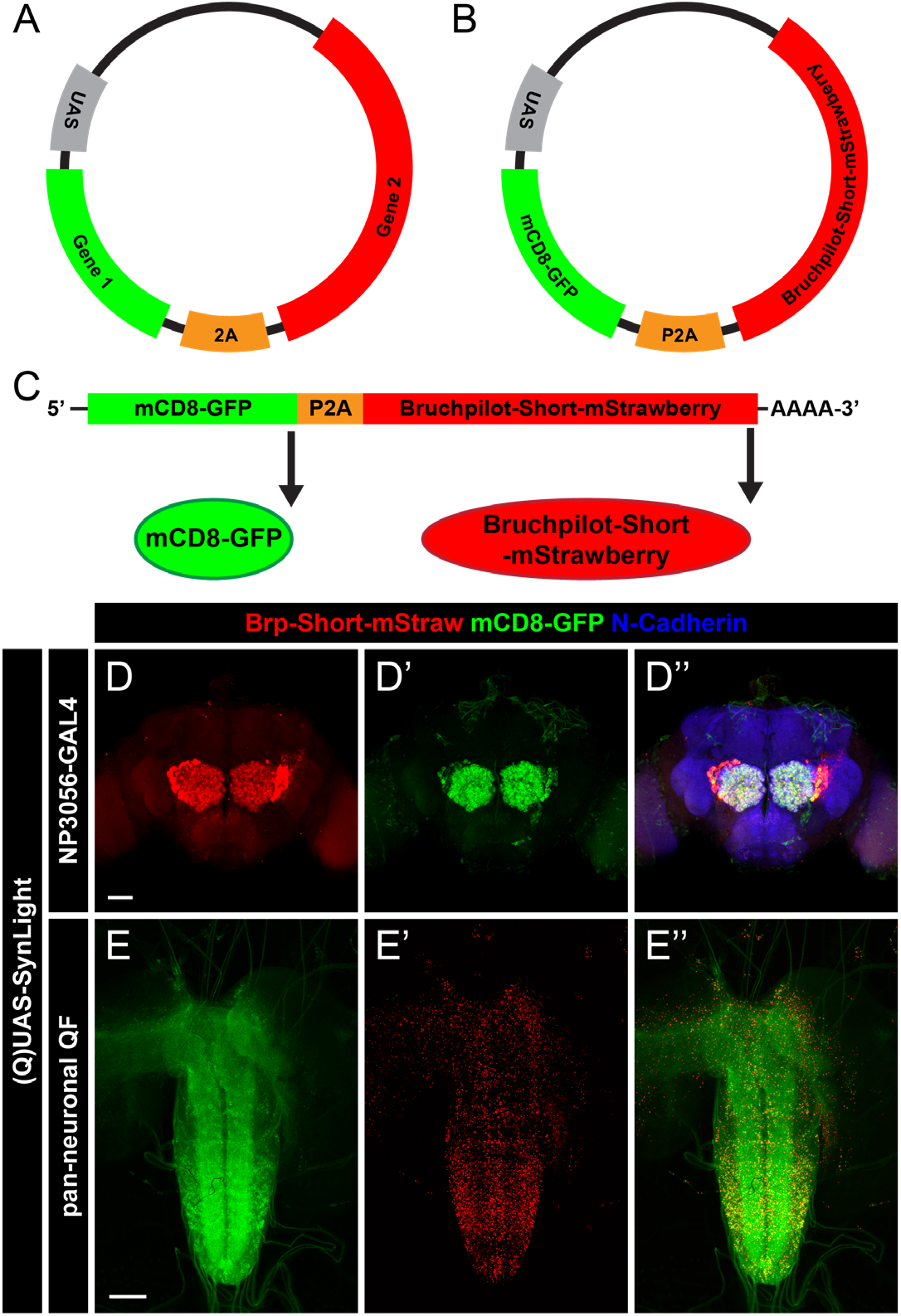
Strategy for generating SynLight, a single transgene that expresses both membrane-tagged GFP and mStrawberry-tagged Bruchpilot-Short. **A**, Diagram of an example plasmid containing a UAS vector and codon-optimized 2A peptide coding sequence (Daniels et al., 2014). Flanking either side of 2A are multiple cloning sites and restriction sites that facilitate insertion of two or more genes of interest. **B**, Diagram of the SynLight plasmid. Using restriction enzymes, the mCD8-GFP coding sequence was inserted preceding the P2A coding sequence and then the Bruchpilot-Short-mStrawberry coding sequence is inserted following the P2A sequence, keeping all sequences in frame. **C**, Diagram of Syn-Light mRNA, showing two separate proteins being produced from a single mRNA sequence. **D-D”**, Representative maximum projection confocal image stacks of multiglomerular LNs of the adult brain expressing Syn-Light and stained with antibodies against mStraw (red), GFP (green), and N-Cadherin (blue). These images show overlapping, yet distinctly different subcellular localization of Brp-Short-mStraw and mCD8-GFP. **E-E”**, Representative maximum projection confocal image of the third instar ventral nerve cord expressing SynLight, showing separate endogenous expression of Brp-Short-mStraw and mCD8-GFP via the native fluorescence from the mStrawberry and GFP fluorophores. Scale bars = 40 μm (D); 80 μm (E).

### Quantification of synapses and neuronal morphology in antennal lobe neurons using SynLight

Both Bruchpilot-Short and mCD8-GFP have enabled quantitative analyses of synaptic organization at peripheral and central synapses, resulting in established measurements of synapse number and neurite volume, especially in the olfactory system (Aimino et al., 2023; Christiansen et al., 2011; Kremer et al., 2010; Mosca et al., 2017; Mosca & Luo, 2014). As such, there is a rich history of control data against which we can benchmark SynLight performance. To demonstrate the utility of SynLight for making quantitative measurements of synapse organization and density, we first turned to olfactory receptor neurons (ORNs) in the *Drosophila* antennal lobe. In the antennal lobe, ORNs, projection neurons (PNs), and local interneurons (LNs) are the three major neuron types that contribute to the sensation and subsequent relay of olfactory information to higher order brain structures such as the mushroom bodies and the lateral horn (Jefferis et al., 2001; Tanaka et al., 2009; Vosshall et al., 2000). ORNs, PNs, and LNs each project to the roughly 50 glomeruli that comprise the antennal lobe, which are subdivided based on the type of olfactory information they receive, and form synapses with each other to create functional circuits (Grabe & Sachse, 2018; Hallem & Carlson, 2006; Jefferis et al., 2007; Suh 2004).

We first examined ORNs of the VA1lm glomerulus (Figure 2A) using Or47b-GAL4 (Vosshall et al., 2000) and compared expression of independent Brp-Short-mStrawberry and membrane-bound GFP transgenes (Figure 2B-C”) to SynLight (Figure 2D-E”) following immunohistochemical staining for Brp-Short-mStraw and mCD8-GFP. Qualitatively, expression patterns and subcellular localization of SynLight versus the mCD8-GFP / Brp-Short-mStraw from independent transgenes were indistinguishable from one another regardless of genotype. Subsequently, for each genotype, we quantified Bruchpilot-Short puncta and neurite volume (as represented by mCD8-GFP staining) and found that Brp-Short puncta number (Figure 2F), neurite volume (Figure 2G), and synapse density (Figure 2H) in VA1lm ORNs were not significantly different between flies expressing SynLight and those expressing Brp-Short-mStrawberry and mCD8-GFP independently. This indicates that SynLight accurately recapitulates independent Brp-Short and mCD8-GFP expression both qualitatively and quantitatively. Moreover, SynLight expression does not interfere with synaptic organization or development of individual neuron types, as the mature synapse number and volume is unaltered when compared to published data (Aimino et al., 2023; Mosca et al., 2017; Mosca & Luo, 2014). Therefore, SynLight is a viable strategy for quantitatively assessing synaptic organization with fewer genetic transgenes.

**Figure 2.**
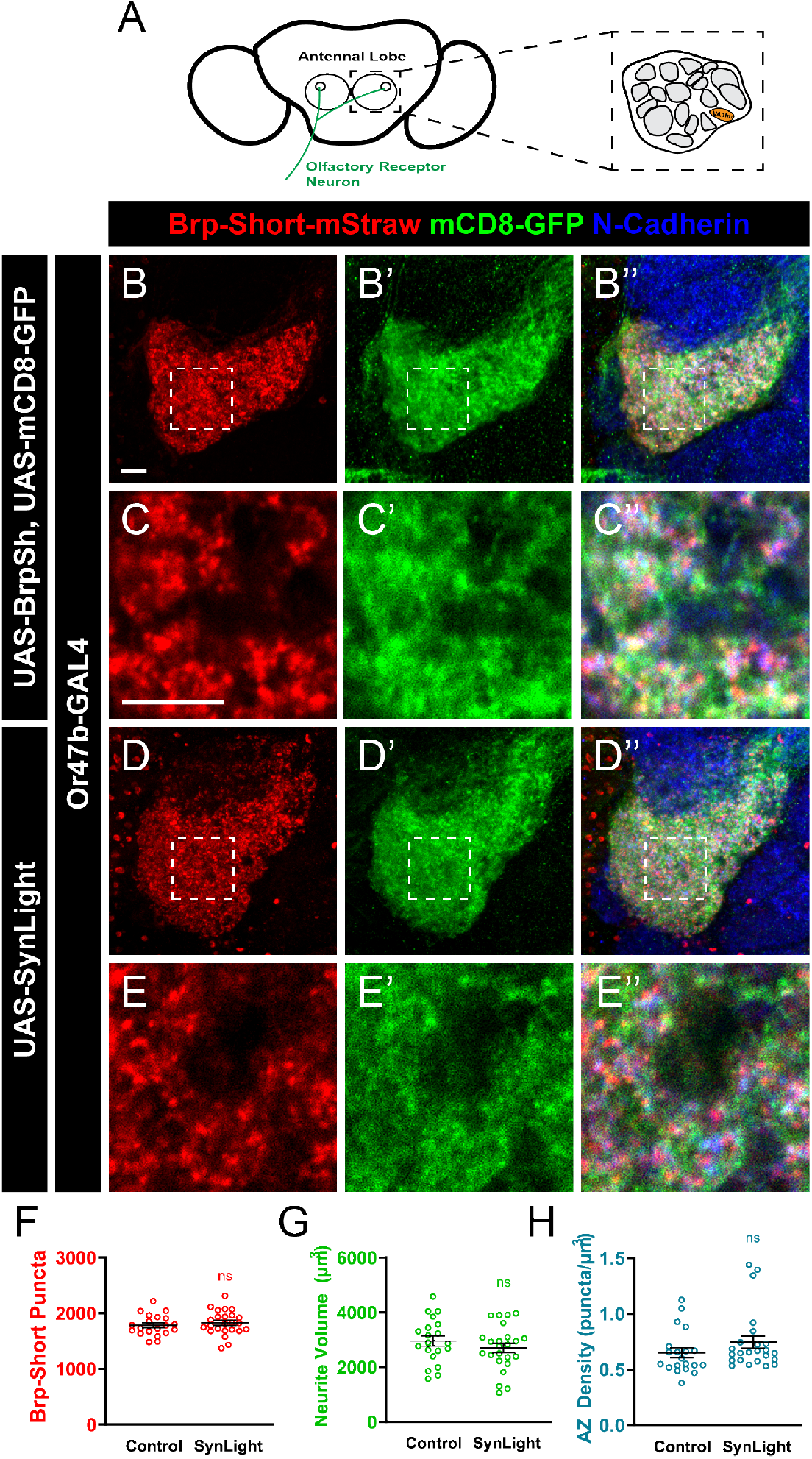
SynLight expression does not affect synapse number in olfactory neurons. **A**, Diagram of the Drosophila antennal lobes showing ORNs (green) of the VA1lm glomerulus (orange). **B-B”**, Representative confocal image stacks of 10-day old male adult VA1lm ORNs expressing Brp-Short-mStraw and membrane-tagged GFP separately and stained with antibodies against mStraw (red), GFP (green), and N-Cadherin (blue). **C-C”**, High-magnification, single optical image section of ORNs from inset in B showing co-localization, but not complete overlap, of synaptic labels. **D-D”**, Representative confocal image stacks of 10-day old male adult VA1lm ORNs expressing SynLight and stained with antibodies as in B. **E-E”**, High-magnification, single optical image section from inset in D also showing co-localization, but not complete overlap, consistent with subcellular localization and suggesting P2A-mediated cleavage is occurring successfully. **F-H**, Quantification of Brp-Short-mStraw puncta number (F), membrane GFP volume (G), and synapse density (H) for adult male VA1lm ORNs expressing either SynLight or Brp-Short-mStraw and membrane-tagged GFP separately. Brp-Short puncta number, neurite volume, and synapse density obtained using SynLight are not significantly different from using Brp-Short-mStraw and membrane-GFP separately. For each genotype, n ≥ 20 glomeruli from 10 brains. n.s. = not significant. Scale bars = 5 μm.

Having established that SynLight is robustly expressed in antennal lobe VA1lm ORNs without affecting synaptic organization, we next expanded our analysis by driving SynLight expression in multiple cell types of the olfactory system (Figure 3A). When driven in a different population of antennal lobe ORNs using Or67d-GAL4 (DA1 ORNs, Stockinger et al., 2005) or Or67d-QF (Liang et al., 2013), we saw robust labeling of ORN active zones and neurites (Figure 3B-C”). We also examined SynLight expression in other antennal lobe neurons beyond ORNs: we used Mz19-GAL4 (Jefferis et al., 2003) and NP3056-GAL4 (Y. H. Chou et al., 2010) to drive SynLight expression in DA1 PNs (Figure 3D-D”) and multiglomerular LNs of DA1 (Figure 3E-E”), respectively. As with DA1 ORNs, we found that SynLight labels active zones and neurites in both classes of neurons and that the labeling is consistent with previous results from the same drivers (Aimino et al., 2023; Mosca & Luo, 2014). Taken together, SynLight expression is evident regardless of the olfactory neuron class in which it is expressed or via which binary expression system it is driven, further demonstrating its utility as a tool for studying synapse formation and organization.

**Figure 3.**
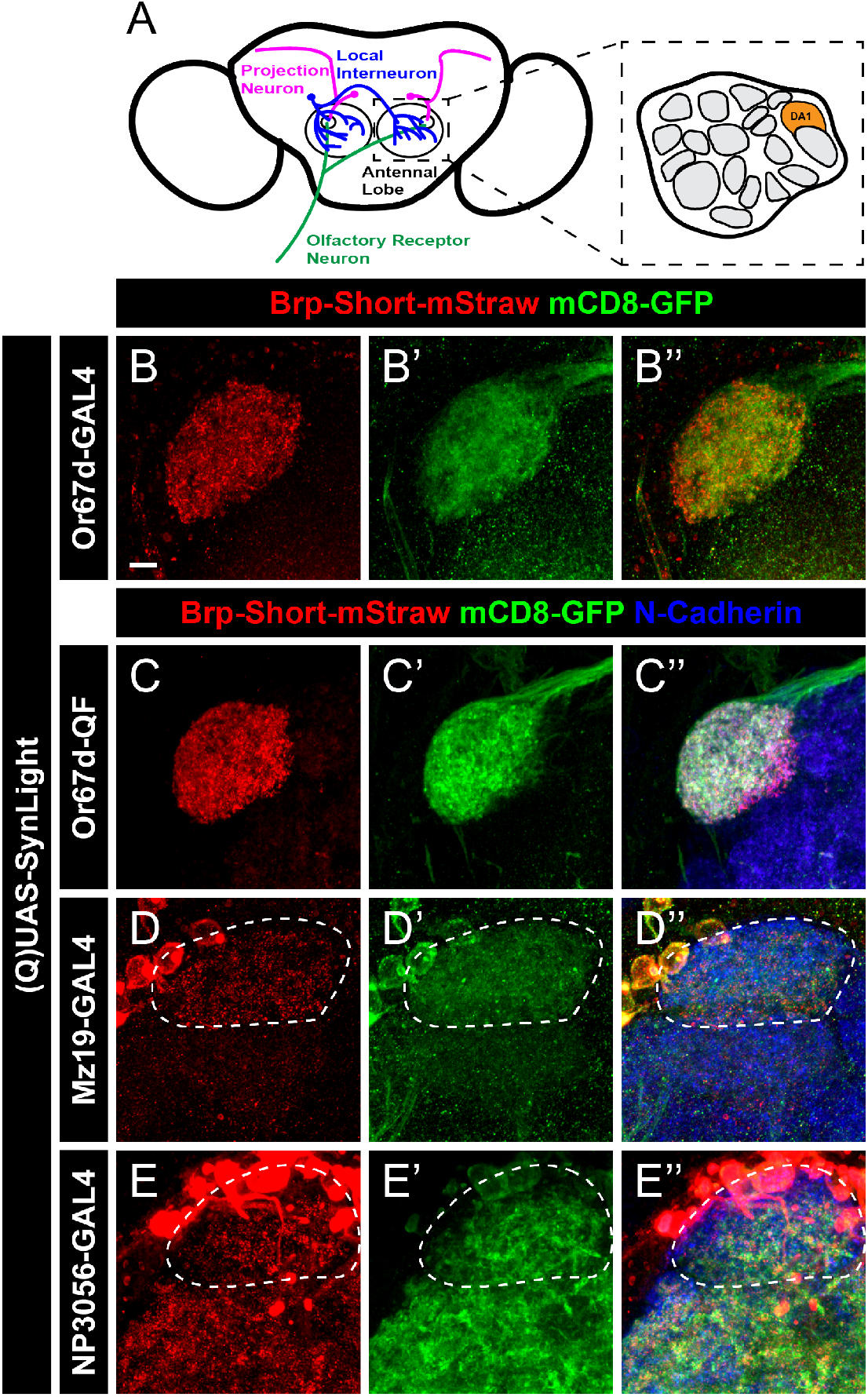
SynLight labels presynaptic active zones and neuronal membranes in multiple cell types of the olfactory system. A, Diagram of the Drosophila antennal lobes showing ORNs (green), PNs (magenta), and multiglomerular LNs (blue) of the DA1 glomerulus (orange). B-C”, Representative confocal image maximum projections of male adult DA1 ORNs expressing SynLight via a GAL4 (B) or QF (C) driver and stained with antibodies against mStraw (red), GFP (green), and N-Cadherin (blue). D-E”, Representative confocal image maximum projections of male adult PNs (D) and multiglomerular LNs (E) of the DA1 glomerulus expressing SynLight and stained with antibodies as in B-C. Scale bar = 5 μm.

### SynLight labels presynaptic connections in neurons of the visual system

To expand our study of SynLight expression beyond the olfactory system, we next examined the fly visual system. Both the anatomy and organization of the fly visual system has been well characterized (Choi et al., 2021; Scheffer et al., 2020; Takemura et al., 2013, 2015; Yang & Clandinin, 2018) and the fly visual system represents an excellent model for studying synaptic development and organization (Clandinin & Zipursky, 2002) as well as visual processing (Yang & Clandinin, 2018). Further, tagged versions of Bruchpilot have been used extensively to characterize both the cellular events underlying, and the molecular mechanisms supporting, synaptic plasticity in the visual system (Araki et al., 2020; Berger-Müller et al., 2013; Chen et al., 2014; Duhart & Mosca, 2022; Kawamura et al., 2020; Osaka et al., 2023; Shimozono et al., 2019; Sugie et al., 2015), highlighting the utility of Brp-based labeling tools in understanding visual biology. To determine if SynLight could be used to concurrently label neuronal membranes and active zones in the visual system, we drove UAS-SynLight in the visual system using two different GAL4 drivers, Dpr Interacting Protein-γ (DIP-γ)-GAL4 (Carrillo et al., 2015) and 27B03-GAL4 (Jenett et al., 2012) and examined both Brp-Short puncta and GFP-tagged neuronal membranes. DIP-γ-GAL4 labels Dm8 neurons in layer M6 of the medulla (Figure 4A-A”) while 27B03-GAL4 drives expression in neurons of the optic lobe (Figure 4B-B”). In both cases, we observed robust expression of SynLight and labeling consistent with release sites (via Brp-Short-mStraw) and general neuronal processes (via mCD8-GFP), indicating the efficacy and applicability of the SynLight construct beyond the olfactory system. The Dm8 neurons labeled by DIP-γ-GAL4 are postsynaptic to R7 photoreceptor neurons and form a connection analogous to the ORN to PN synapses in the antennal lobe (Figure 4C; Takemura et al., 2013). Processes of Dm8 neurons also form synaptic contacts onto Tm5c neurons, comprising a circuit that mediates UV preference (Karuppudurai et al., 2014). Both connections (R7-->Dm8 and Dm8-->TM5c) form within the M6 layer of the medulla, suggesting that presynaptic R7 terminals should localize near (but not overlap with) Dm8 presynaptic terminals. We reasoned that concurrent labeling of R7 and Dm8 terminals would result in presynaptic staining of both neuron classes and that their respective presynaptic sites would be found in close proximity to one another within layer M6 of the medulla. To do so, we drove expression of SynLight in Dm8 neurons and co-stained the optic lobes with antibodies to Chaoptin, a marker for R7 photoreceptor cells (Krantz & Zipursky, 1990). Indeed, when we specifically examined the M6 layer, we found that Dm8 Brp-Short puncta and R7 photoreceptor Chaoptin are present in similar regions of the optic lobe (Figure 4D-D”). Furthermore, Dm8 Brp-Short puncta and R7 Chaoptin signals do not overlap, but are instead adjacent to one another as predicted (Figure 4D”’). Thus, SynLight expression can recapitulate expected patterns of synaptic organization in the visual system, indicating its utility as a synaptic label. Taken together with our findings from the olfactory system (Figures 2-3), these data show that SynLight is a robust, reliable tool for concurrent labeling of synaptic active zones and general neuronal processes in multiple central nervous system populations.

**Figure 4.**
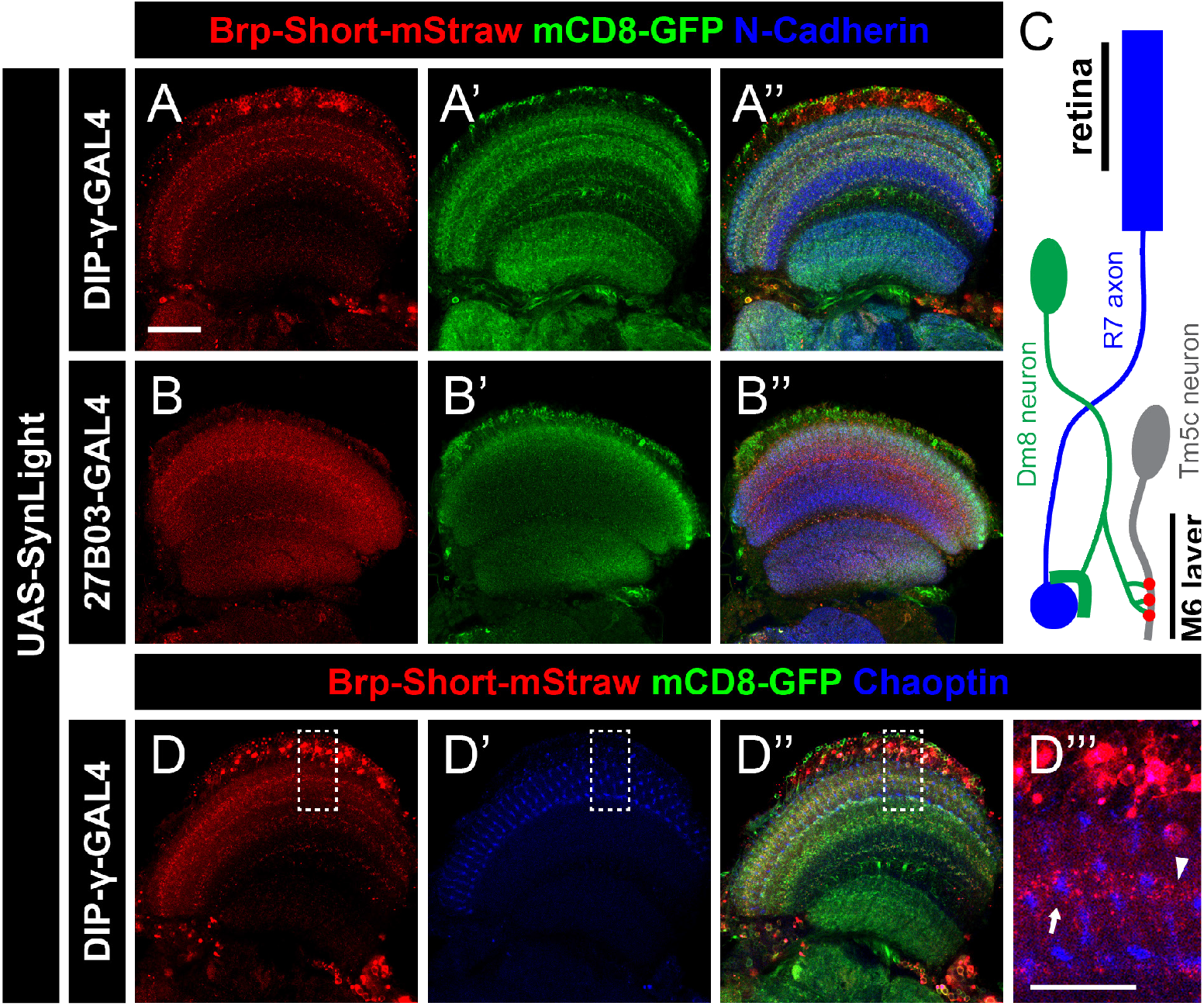
SynLight labels presynaptic active zones and neuronal membranes in neurons of the visual system. A-B”, Representative single confocal image sections of male adult brains expressing SynLight using DIP-γ-GAL4 to label Dm8 neurons (A) or 27B03-GAL4 to label optic lobe neurons (B) and stained with antibodies against mStraw (red), GFP (green), and N-Cadherin (blue). C, Schematic showing the connections between R7 photoreceptor axons (blue), Dm8 neurons (green), and Tm5c neurons (gray). R7 axons project from the retina and synapse onto the dendrites of Dm8 neurons. Dm8 neurons subsequently form synapses with Tm5c neurons, forwarding the visual information received from R7 axons. The presynaptic active zones of Dm8 neurons (red) and axon terminals of R7 cells (blue) are both found in the M6 layer of the medulla. D-D”, Representative single confocal image sections of male adult brains expressing SynLight in Dm8 neurons and stained with antibodies against mStraw (red), GFP (green), and Chaoptin (blue). D”’, Single, high-magnification image section from insets (dashed boxes, D) showing mStraw and Chaoptin costaining. Arrow indicates region of Brp-Short and Chaoptin in close proximity while arrowhead indicates a region with only Brp-Short. Scale bars = 20 μm (A); 10 μm (D”’).

### SynLight accurately labels and quantifies active zones at neuromuscular synapses

To explore the utility of SynLight beyond the central nervous system, we next turned to peripheral neuromuscular junction (NMJ) synapses. NMJ synapses are highly stereotyped and are a long-studied, powerful system for uncovering active zone biology (V. T. Chou et al., 2020a; Landgraf & Thor, 2006; Menon et al., 2013) making them an optimal synapse for examining SynLight expression and quantification. We first expressed SynLight pan-neuronally via elavC155-GAL4 (Lin & Goodman, 1994) and observed robust labeling of both general membranes (via mCD8-GFP) and active zones (via Brp-Short-mStraw) at NMJs (Figure 5B-B”) that was absent from non-expressing controls (Figure 5A-A”). Consistently, mStraw-positive Brp-Short puncta labeled by SynLight overlapped with endogenous Bruchpilot antibody staining (Figure 5C-C”), suggesting that SynLight labeling accurately revealed endogenous active zones. We further observed similar Brp-Short-mStraw and mCD8-GFP expression and localization with QUAS-SynLight (Figure 5D-D”) expression via n-syb-QF (Riabinina et al., 2015), indicating that multiple binary expression system versions of SynLight provide robust labels. Taken together, this indicates that SynLight is effectively and accurately expressed at NMJ synapses in separable pools reflecting membranes and release sites.

**Figure 5.**
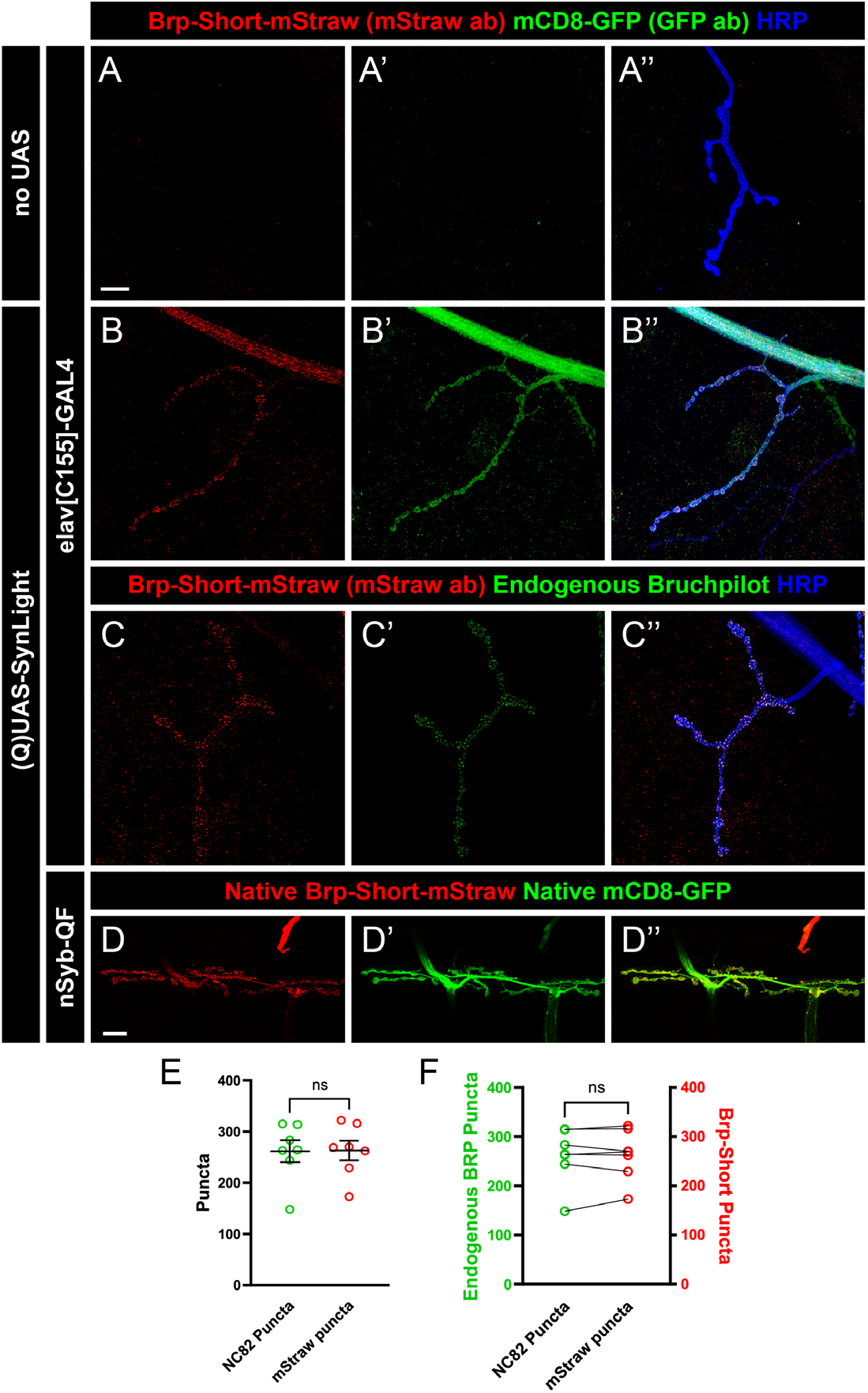
SynLight labels the larval neuromuscular junction and does not alter synapse formation. A-B”, Representative confocal image maximum projections of muscle 4 NMJs in control (A) or SynLight-expressing (B) wandering third instar larvae stained with antibodies against mStraw (red), GFP (green), and HRP (blue). The negative control lacking SynLight shows no mStraw or GFP immunoreactivity while pan-neuronal SynLight expression shows clear visibility of both markers. C-C”, Representative confocal image maximum projections of a muscle 4 NMJ expressing pan-neuronal SynLight and stained for antibodies against mStraw (red), NC82 (green), and HRP (blue). D-D”, Representative confocal image maximum projections of muscle 6/7 NMJs expressing SynLight showing endogenous expression of Brp-Short-mStraw (red) and mCD8-GFP (green). E, Quantification of active zone puncta visualized by antibody staining of endogenous Bruchpilot (via monoclonal antibody NC82) or expression of Brp-Short via SynLight from C. There is no significant difference between Brp-Short-positive and NC82-positive puncta. F, Quantification of active zone puncta from C when separated into each individual NMJ corroborates there is no significant difference between Brp-Short-positive and NC82-positive puncta number. For each experimental group, n ≥ 7 NMJs. n.s. = not significant. Scale bars = 15 μm (A); 20 μm (D).

Having established that SynLight accurately localizes to NMJ synapses and membranes, we next assessed SynLight as a quantitative tool for active zone puncta. NMJ terminals have a characteristic number of active zone puncta when stained with antibodies to endogenous Bruchpilot, highlighting this metric as a reliable quantitative measurement of synaptic growth (Collins & DiAntonio, 2007; Daniels et al., 2008; Wairkar et al., 2008). To determine if SynLight could be reliably used to quantify active zones, we counted Bruchpilot-Short puncta at muscle 4 NMJ terminals and compared the data to counts of puncta recognized by the monoclonal antibody NC82 to endogenous Bruchpilot (Laissue et al., 1999; Wagh et al., 2006). There was no significant difference in the average puncta number visualized by mStraw (via SynLight) or NC82 (monoclonal antibody to Brp) staining (Figure 5E), suggesting that SynLight could accurately quantify Brp-positive endogenous active zone number. Moreover, when we compared Brp-Short-mStraw vs. NC82 puncta counts for each NMJ, we observed no significant difference at the individual NMJ level (Figure 5F), demonstrating accurate and congruent reporting. In all, the data indicates that SynLight accurately reports NMJ synaptic organization both qualitatively and quantitatively. Combined, our findings establish that SynLight functions as a robust synaptic label at both peripheral and central synapses in *Drosophila*.

## DISCUSSION

As technologies improve, making novel manipulations of, and labeling in, the nervous system possible, there is a growing need to incorporate more genetic components into experiments. Experiments in model organisms especially often have at least three transgenes (V. T. Chou et al., 2020b; Duhart & Mosca, 2022; Venken, Schulze, et al., 2011) for even basic experiments: a genetic driver (e.g. GAL4, QF, lexA, Cre), a reporter (GFP, synaptic labels, receptor labels), and an effector (e.g., an optogenetic regulator of activity, toxin, endocytosis blocker, kinase activity regulator). Multiple challenges exist, however, with such experiments. First, each transgene must be accounted for in genetic crosses to obtain experimental animals, leading to complex crosses where it is increasingly challenging to obtain “correct” progeny based on Mendelian ratios and unanticipated lethality. Second, genetic driver strength can be diluted by multiple transgenes (Brand & Perrimon, 1993), leading to increased variability of expression and / or reduced efficacy of expressed transgenes. Finally, space constraints (from chromosome number or limits on viral DNA payload) can limit the number of genetic transgenes that can be present in the final experimental animal. Though some approaches like recombination of multiple transgenes onto the same chromosome can increase available genetic space for other transgenes and alleviate some of these concerns, recombinants do not reduce the total number of transgenes and the “genetic load” of the system persists. To begin to address some of these concerns, we developed a new strategy, SynLight, that uses the viral P2A ribosomal skipping peptide (Daniels et al., 2014; Diao & White, 2012) to produce a single transgene that expresses both the membrane label mCD8-GFP (T. Lee et al., 1999) and the active zone label Bruchpilot-Short-mStrawberry (Fouquet et al., 2009; Mosca & Luo, 2014). We demonstrate that this strategy is effective in multiple central and peripheral neurons and is quantitatively similar to synaptic measurements using independent mCD8-GFP or Bruchpilot-Short expression alone (Aimino et al., 2023; Mosca & Luo, 2014). Using SynLight in either CNS or PNS experiments will enable more complex studies in vivo without sacrificing the number of labels possible.

To develop a construct for use in *Drosophila* that encodes both a presynaptic active zone marker as well as a neuronal membrane tag from a single sequence, we incorporated the P2A peptide, a ribosomal skipping sequence (Luke et al., 2009). This virus-derived peptide sequence mediates a skipping event during translation that enables the production of both proteins from a single mRNA (Daniels et al., 2014; Kim et al., 2011; Tang et al., 2009). We incorporated the established fly active zone label Bruchpilot-Short (Aimino et al., 2023; Christiansen et al., 2011; Duhart & Mosca, 2022; Fouquet et al., 2009; Kittel et al., 2006; Kremer et al., 2010; Mosca & Luo, 2014; Wagh et al., 2006) and the general membrane marker mCD8-GFP (T. Lee et al., 1999) to produce a single SynLight transgene that concurrently labels all neuronal membranes via mCD8-GFP and mature active zones via Brp-Short. We established multiple SynLight transgenic constructs (Figure 1) on the 2nd and 3rd chromosomes for GAL4/UAS expression (Brand & Perrimon, 1993) and on the 3rd chromosome for QF/QUAS expression (Potter et al., 2010). We further demonstrated that SynLight is expressed with high fidelity in multiple classes of *Drosophila* CNS neurons, including those of the olfactory (Figures 2-3) and visual (Figure 4) systems. Further, the two products, mCD8-GFP and Bruchpilot-Short-mStraw, are readily separable in all neurons and, when quantified, produce similar results to established data and to expression of individual analogous constructs alone (Figure 2). Thus, SynLight is effective for quantifying synapse density with only one construct, whereas previous experiments required two independent transgenes. We also demonstrated similar utility for SynLight at peripheral NMJ synapses. Not only is expression robust and labeling evident (Figure 5) for membranes and active zones; measurements with SynLight accurately recapitulate data obtained from established antibodies to endogenous Bruchpilot (Laissue et al., 1999; Wagh et al., 2006). In all, SynLight accurately labels multiple subcellular structures via only one transgene.

Tools like SynLight will allow greatly increased utility within the fly nervous system. This will promote not only more complex and nuanced questions, but also reduce experimental work. For example, to determine whether reduction of the function of a single gene influences synaptic density, five transgenes would optimally be required: a genetic GAL4 / QF / lexA driver, an RNAi transgene to reduce specific gene function, Dcr2 to increase RNAi efficacy (Dietzl et al., 2007), mCD8-GFP (or an equivalent membrane marker) to measure neurite volume, and Brp-Short (or an equivalent active zone label) to quantify release sites. Not only is this a genetically complex experiment, it may also reduce expression when a driver is used to express four independent transgenes. While the expression level tendered by strong drivers will enable the experiment, many circumstances will result in either reduced expression of the labels, and / or reduced efficacy of the RNAi, leading to difficulty in interpreting the results. Previous approaches (Aimino et al., 2023; Mosca et al., 2017) have expressed the mCD8-GFP and Bruchpilot-Short transgenes in separate experiments, but as the measurements are then not taken from the same animal, synaptic density is not directly calculable. SynLight circumvents that issue by using only a single transgene to express both labels, thus increasing the utility of available experiments. Beyond simple perturbation experiments using a single class of neurons, SynLight also enables more complex, transsynaptic questions. When multiple binary expression systems are needed to label and manipulate different neuronal populations, as with pre- and postsynaptic neurons (Mosca & Luo, 2014; Parisi et al., 2023), the required experiments must be carefully designed with limits on the ensuing number of transgenes (since multiple genetic drivers now contribute to the genetic load). In this case, one expression system would drive the expression of synaptic labels in one neuronal population while another expression system would drive an effector transgene in a second neuronal population. Employing a construct such as SynLight, which codes for multiple proteins from a single sequence, reduces the genetic load on the system and makes it easier to produce the correct experimental fly stocks in the absence of genetic dilution. Additionally, this transgene can be recombined with other transgenes, further simplifying the creation of a desired stock. Finally, our use of SynLight presents further proof-of-principle of the utility of 2A peptides in vivo in *Drosophila*. Prior work established transgenes containing a Ca^2+^ reporter like GCaMP and a membrane label (Daniels et al., 2014). Use of T2A to produce GAL4 expressing constructs at the end of a protein reporter or endogenous protein have also greatly enhanced neuronal circuit study (Diao & White, 2012; Kondo et al., 2020; P. T. Lee et al., 2018). By including two different reporters for membranes and active zones, this greatly increases the number and kinds of experiments possible. Future versions can pair effectors and labels (like Bruchpilot-Short and an activity-altering construct) or even enzymes and labels (like a FLPase and Bruchpilot-Short) as needed to design different kinds of experiments. Overall, SynLight enables the high-resolution visualization of presynaptic active zones as well as neuronal membranes in vivo via a single transgene, thus reducing the genetic load on the system. We anticipate that this new approach be applied throughout the central and peripheral nervous system to answer more complicated questions about circuit biology, neurodevelopment, and synaptic organization.

## ACKNOWLEDGMENTS

We would like to thank all members of the Mosca lab for their constant support, input, and critical discussion of the project. We thank Dr. Kristen Davis, Alison DePew, and S. Zosimus for comments on the manuscript. We thank the Developmental Studies Hybridoma Bank (created by the NICHD of the NIH and maintained at The University of Iowa, Department of Biology, Iowa City, IA 52242) for antibodies. Stocks obtained from the Bloomington *Drosophila* Stock Center (NIH P40OD018537) were used in this study and we thank them for their service to the community.

## AUTHOR CONTRIBUTIONS

M.A.A. and T.J.M. designed the project; M.A.A., J.H., M.J.P., and J.C.D. performed experiments; M.A.A., M.J.P., and J.C.D. produced reagents; M.A.A. and J.H. analyzed the data. M.A.A., J.H., M.J.P., J.C.D., and T.J.M. wrote and edited the manuscript.

## DATA AVAILABILITY

All fly lines are available upon request. The final construct sequences for all variants of SynLight are also available upon request.

